# Dynamics and Energetics of Bottlenose Dolphins (*Tursiops truncatus*) Fluke-and-Glide Gait

**DOI:** 10.1101/2022.04.19.488827

**Authors:** Ding Zhang, Yifan Wang, Joaquin Gabaldon, Lisa K. Lauderdale, Lance J. Miller, Kira Barton, Kenneth Alex Shorter

## Abstract

Intermittent locomotion composed of periods of active flapping/stroking followed by inactive gliding has been observed with species that inhabit both aerial and marine environments. However, studies on the energetic benefits of a fluke-and-glide (FG) gait during horizontal locomotion are limited for dolphins. This work presents a physics-based model of FG gait and analysis of the associated costs of transport of bottlenose dolphins (*Tursiops truncatus*). New estimates of gliding drag coefficients for the model were estimated experimentally from free-swimming bottlenose dolphins. The data-driven approach used kinematic measurement from 84 hours of biologging tag data collected from 3 animals to estimate the coefficients. A set of 532 qualified gliding events were automatically extracted for gliding drag coefficient estimation, and an additional 783 FG bouts were parameterized and used to inform the model-based dynamic analysis. Experimental results indicate that FG gait was preferred at speeds around 2.2 - 2.7 m/s. Observed FG bouts had an average duty factor of 0.45 and gliding duration of 5 sec. The average associated metabolic cost of transport (COT) and mechanical cost of transport (MECOT) of FG gait are 2.53 and 0.35 J · m^*−*1^ · kg^*−*1^ at the preferred speeds. This corresponded to an 18.9% and 27.1% reduction in cost when compared to modeled continuous fluking gait at the same reference speed. Average thrust was positively correlated with fluking frequency and amplitude as animals accelerated during the FG bouts. While fluking frequency and amplitude were negatively correlated for a given thrust range. These results support the supposition that FG gait enhances the horizontal swimming efficiency of bottlenose dolphins and provides new dynamical insights into the gait of these animals.

## INTRODUCTION

Efficient locomotion is essential for animals traveling long distances, and various morphological or behavioral strategies have been observed to this end (Feldkamp (1987); Au and Weihs (1980); Hui (1989); Fish and Hui (1991); Weimerskirch et al. (2001); Williams et al. (1992); Weihs (2004)). Intermittent locomotion (Kramer and McLaughlin (2001)), where the animal employs a gait composed of periods of active flapping/stroking/bursting and inactive gliding/coasting, is a convergent evolutionary trend in both aerial and marine animals (Gleiss et al. (2011)) that has been observed with birds and fishes (Tobalske (2001); Rayner et al. (2001); Watanuki et al. (2005); Weihs (1973); Fish et al. (1991); Müller et al. (2000)). While the energetic benefits of intermittent locomotion have been studied for certain fishes from various perspectives (Weihs (1973); Fish et al. (1991); Müller et al. (2000); Lighthill (1971); Akoz and Moored (2018); Weihs (1974); Videler and Weihs (1982); Ashraf et al. (2020)), little work has been done with dolphins. Because dolphins are excellent swimmers, and burst-and-coast swimming is not always energetically beneficial in fish (Ashraf et al. (2020)), it is important to investigate the energetic trade-offs of intermittent locomotion. For cetaceans, published work has focused on how animals extend dive duration and conserve energy by taking advantage of buoyancy changes while gliding during descent or ascent in a dive (Williams (2001); Williams et al. (2000); Skrovan et al. (1999)). However, a model-based investigation of the energetic benefits of intermittent gait during horizontal locomotion, specifically a fluke-and-glide (FG) gait, is lacking for these animals. This work will investigate the following questions: When is FG gait energetically beneficial? Do animal speeds observed during FG gait have a reduced energetic cost than continuous fluking? Are the relationships between kinematic parameters (e.g., fluking frequency vs. speed) observed during steady-state swimming held for FG gait?

Here we are interested in comparing the energetic cost between continuous and FG gait and the associated dynamical conditions during intermittent locomotion. Researchers have investigated the metabolic effort of dolphins during prescribed trials by estimating energetic cost using respirometry and blood lactate measurements. The prescribed trials have included boat following at a constant speed (Williams et al. (1992)); static force generation (Williams et al. (1993)); and point-to-point swimming at different speeds and levels of effort (Yazdi et al. (1999); Williams et al. (2017); van der Hoop et al. (2018; 2014)). However, gait during self-selected swimming spans a continuum that includes continuous and intermittent fluking. It is not easy to train the animals to swim with a prescribed gait pattern. As a result, experimental estimates of cost from respiratory or blood measurements can be limited because it is difficult to decouple the mixed effects of different motion modes, particularly when these modes are combined dynamically during the trials. Thus, an explicit dynamic model is particularly useful for investigating a combined gait composed of multiple motion modes with varying ratios.

A model-based evaluation of energetic cost requires an accurate estimate of the hydrodynamic drag experienced by the animal during both fluking and gliding. Various methods have been used to obtain such estimations for marine animals, including theoretical computations (Lighthill (1971); Akoz and Moored (2018); Hertel (1966)), computational fluid dynamic simulations (Zhang et al. (2020); van der Hoop et al. (2018; 2014)), towing experiments (Purves et al. (1975)), camera assisted experiments (Zhang et al. (2020); Noren et al. (2011); Feldkamp (1987); Skrovan et al. (1999)), and field biologging tag deployments where speed can be derived from the data to estimate drag (Miller et al. (2004)). However, estimates of gliding drag over a range of speeds for bottlenose dolphins (*Tursiops truncatus*) are currently lacking. In this work, we estimate gliding drag coefficients for dolphins experimentally by fitting a drag model to speed measurements during gliding (Lang (1975); Bilo and Nachtigall (1980); Stelle et al. (2000); McHenry and Lauder (2005); Zhang et al. (2020)). In contrast to studies that use video data from structured trials to derive gliding speed profiles, we took a data-driven approach to extract the information from self-selected swimming data collected using biologging tags.

Using these new coefficients, we characterize the FG gait of bottlenose dolphins during self-selected swimming. Specifically, the gliding drag coefficient, gliding duration, reference speed, and duty factor during observed FG gait are used in a dynamic model to estimate energetic cost. We model the FG gait of bottlenose dolphins as a two-phase process and quantify the associated thrust force, metabolic energy cost of transport (COT), and mechanical energy cost of transport (MECOT) of the animal over a range of swimming gaits. Custom biologging tags were used to directly measure forward speed, three-axis acceleration, orientation, and depth of the animal during self-selected swimming. Swimming gaits were parameterized using tag data by calculating duty factor and glide duration from animal kinematics. These swimming characteristics informed the range of input parameters for the FG gait model used to evaluate the COT and MECOT. Finally, fluking frequency and amplitude of the animal during the accelerating fluking phase of the FG gait were extracted from tag data to investigate swimming biomechanics. New information about gliding hydrodynamics, fluking gait parameters, and the model-based estimates of COT and MECOT during horizontal intermittent locomotion provide further insight into the swimming biomechanics of bottlenose dolphins.

## MATERIALS AND METHODS

### Data collection

Three bottlenose dolphins (*Tursiops truncatus*), (TT01, TT02, TT03), from the Brookfield Zoo, Chicago Zoological Society, participated in this study: age (16, 5, and 38 years), mass (153, 150,and 202 kg), body length (242, 226, and 266 cm) and girth (128, 127, and 139 cm). A biologging tag was attached to the animal 20 cm behind the blowhole via four silicone suction cups (Fig. 1-a) by an animal care specialist. The ESTAR Lab (University of Michigan) designed and built the biologging tag with the tag electronics (OpenTag, Loggerhead Instruments, FL) encapsulated inside a 3D printed tag housing. Sensors onboard include a three-axis accelerometer, gyroscope, magnetometer (50 Hz), pressure sensor, temperature sensor, and speed sensor (10 Hz). The speed sensor measures relative fluid speed by counting the revolving frequency of a microturbine on top of the tag, which is then mapped to the animal’s forward speed in water (Gabaldon et al. (2019)). In addition, software from a custom gait analysis toolbox for the tag (Zhang et al. (2021)) was used to calculate orientation and identify gait parameters. The animal swam freely in the main aquarium habitat (33.5 m (length) by 12.2 m (width) by 6.7 m (depth)) during the deployments. The tags were deployed for an average of 2 hours at a time (84 hrs total).

**Fig. 1.**
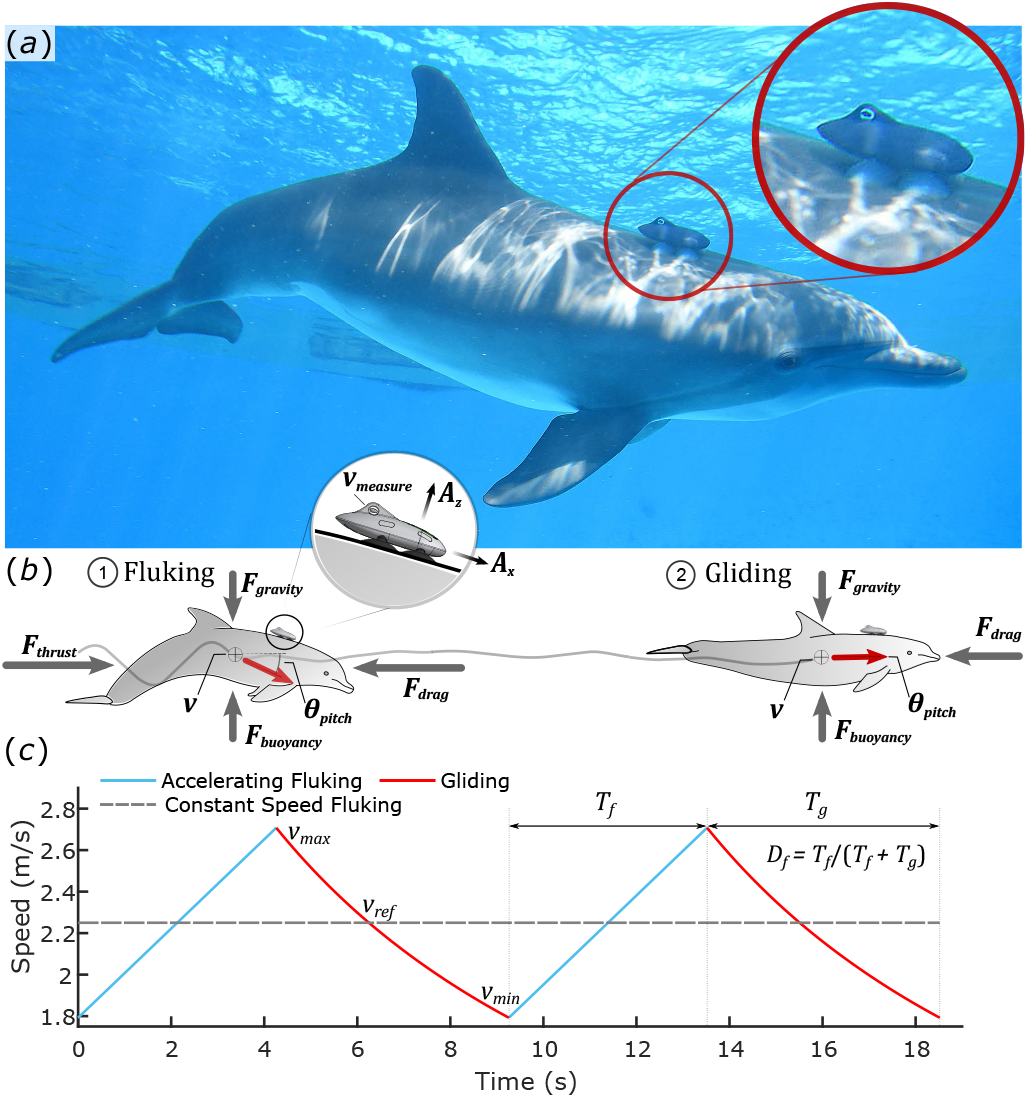
Experiment environment and concepts. (a): A bottlenose dolphin with a biologging tag. (b): Simplified free-body diagrams of an animal fluking and then gliding. (c): Simplified speed model of fluke-and-glide gait and continuous fluking gait over two cycles.

### Dynamics of the fluke-and-glide (FG) gait

Similar to the flapping and gliding model for birds (Sachs (2015)), dolphin fluke-and-glide (FG) gait is modeled as a two-phase process in this work (Fig. 1-c): (1) an accelerating fluking phase with duration *T*_*f*_ and (2) a decelerating gliding phase with duration *T*_*g*_. The maximum and minimum speeds in this process are denoted by *v*_*max*_ and *v*_*min*_, with the constant reference speed:

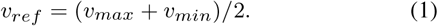

We further simplify the scenario by assuming that the animal is neutrally buoyant and swimming straight (Fig. 1-b). With these assumptions, the thrust force during fluking at time instance *t* can be related to the hydrodynamic drag force *F*_*drag*_ (*t*) and animal’s acceleration *a*(*t*) by:

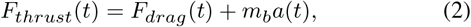

where *m*_*b*_ is the induced virtual mass of the animal (Vogel (1996)). The drag force *F*_*drag*_ (*t*) opposes the direction of motion and is modeled in a conventional way (Fish et al. (2014)) by:

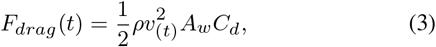

where *ρ* is the density of seawater, *v*(*t*) is the forward speed with respect to water at *t, A*_*w*_ is the wetted surface area of the animal (Fish (1993)), and *C*_*d*_ is the dimensionless drag coefficient for the animal.

The drag coefficient during fluking is denoted as 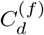 and during gliding as 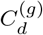. These two parameters are critical for the estimation of drag during FG gait. In this work, we use the 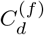 estimated for bottlenose dolphins by Fish et al. (2014), and estimate 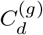 using biologging tag data.

During a glide, there is no thrust force and Eq. (2) reduces to:

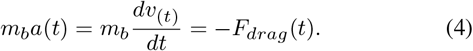

Combining Equations (3) and (4), we obtain the differential equation that describes how speed *v*_(*t*)_ changes during a glide as:

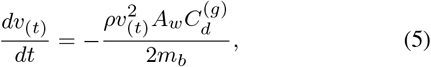

which can be used to solve for speed *v*_(*t*)_ as a function of time *t* and initial speed *v*_(0)_:

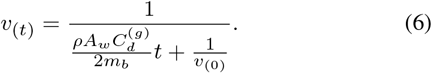

Equation (6) is then used with *measured* speed data during periods of gliding to identify the drag coefficient 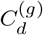, as in Zhang et al. (2020).

Drag coefficients are assumed to be related to Reynolds number (*Re*) by *C*_*d*_ = *bRe*^*c*^ (Fish et al. (2014)), which is linear in the log scale. The Reynolds number for each gliding event is calculated as:

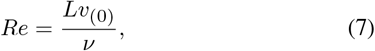

where *L* is the body length of the animal, *v*_(0)_ is the identified initial speed of the glide, and *ν* is the kinematic viscosity of seawater.

### Kinetics of the fluke-and-glide (FG) gait

With the gliding drag coefficient 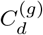 identified, parameters extracted from measured speed data were then used to construct a simplified speed profile (Fig. 1-c) that was used to investigate swimming kinetics with the proposed FG gait model. The speed profile can be uniquely determined given a constant reference speed *v*_*ref*_, a gliding duration *T*_*g*_, and a duty factor *D*_*f*_ which relates *T*_*g*_ and *T*_*f*_ via *D*_*f*_ = *T*_*f*_ */*(*T*_*f*_ + *T*_*g*_). The starting and ending speeds of the glide (i.e. *v*_*max*_ and *v*_*min*_) can be derived from Eq. (6) by providing:

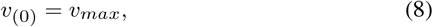

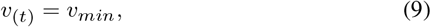

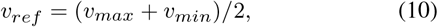

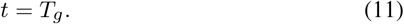

Resulting with:

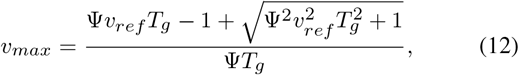

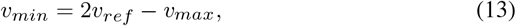

where

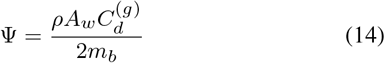

is the coefficient of *t* in Eq. (6). Then, the fluking duration *T*_*f*_ and acceleration *a*(*t*) can be obtained as:

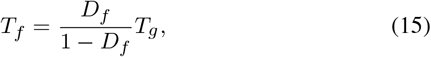

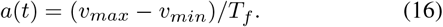

The entire speed profile of a FG gait cycle can be thus represented as:

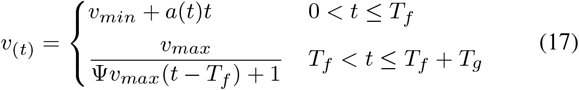

with the two lines corresponding to the speed profile during fluking and gliding, respectively. In the model, *v*(*t*) can be uniquely determined with a triplet of *v*_*ref*_, *T*_*g*_, and *D*_*f*_.

### Cost of transport

Metabolic energy cost of transport (denoted as COT, Schmidt-Nielsen (1972)) characterizes the amount of metabolic or chemical potential energy (*E*, in joules) required to transport one unit of body mass (total *m*, in kilograms) over one unit of distance (total *S*, in meters):

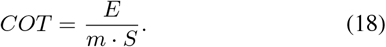

The metabolic energy cost (*E*) can be estimated via measured oxygen consumption, heart rate, and/or plasma lactate concentration during exercise (Williams et al. (1992; 1993; 2017); Yazdi et al. (1999)). Alternatively, we estimated the metabolic energy (*E*) by combining estimated resting metabolic energy (*E*_*res*_) with an estimate of the mechanical work (*W*) produced by the animal during fluking, similar to (Gabaldon et al. (2022)):

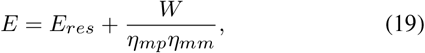

where *η*_*mp*_ = 0.85 is the estimated muscle-to-propulsion conversion efficiency for bottlenose dolphin (Fish (1998)) and *η*_*mm*_ = 0.25 is the estimated mammalian metabolic-to-muscle efficiency (Faraji et al. (2018)). *E*_*res*_ was obtained by integrating an estimated resting metabolic rate (*P*_*res*_) over time (van der Hoop et al. (2014)):

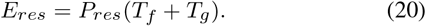

For a bout of FG gait (Fig. 1-c), we calculated the traveled distance *S* and estimated mechanical work *W* via:

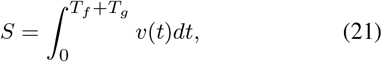

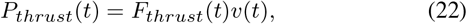

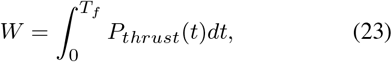

where *F*_*thrust*_(*t*) was obtained from Eq. (2) together with the speed profile *v*(*t*) of the proposed FG model (Eq. (17), Fig. 1-c). *P*_*thrust*_(*t*) represents the thrust power.

In addition to the metabolic cost of transport (COT), we also used mechanical energy cost of transport (MECOT) to quantify the minimum amount of mechanical energy (in joules) required to transport one unit of body weight (1 kilogram) over one unit of distance (1 meter):

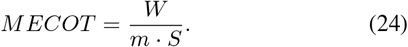

Unlike COT, MECOT only includes the dynamic of the animal.This simplified metric can be used to compare the gait across individuals, potentially other mechanical systems like a bioinspired robotic dolphin (Wu et al. (2019); Conti (2021)).

The FG model, COT, and MECOT were used to investigate how swimming parameters are related to cost during FG gait. Reference swimming speed *v*_*ref*_, gliding duration *T*_*g*_, and duty factor *D*_*f*_ were extracted from the experiment data and used to determine *v*(*t*) (Eq. (17)) as well as other dynamic properties and the cost of transport. Constant speed continuous fluking gait (i.e. *v*_(*t*)_ = *v*_*ref*_, *T*_*g*_ = 0, *D*_*f*_ = 1) was investigated using the model for comparison.

### Glide extraction and FG bout identification

Calibrated accelerometer, gyroscope, and magnetometer data were used to estimate the animal’s orientation (pitch, roll, and yaw) using a gradient-descent-based optimization method in the quaternion space (Madgwick et al. (2011)). When the animals were moving (forward speed *>* 1.0 m/s), three criteria were empirically used to identify gliding events for analysis. First, the glide had to occur at a depth greater than 2.5 body diameters to avoid surface drag (Hertel (1966)). For all animals, a depth greater than 3.5 m satisfied this condition. Second, the animal was not actively swimming or changing direction during the gliding event. The postural configuration and the use of control surfaces (fins and flukes) for maneuvering would affect drag acting on the body. The 1-norm of jerk (i.e. the derivative of the 3-axis measurements from the accelerometer) ||jerk||_1_ was required to be less than 0.5 g/s to exclude active swimming, and restricted angular rates (pitch rate < 20 deg/s, yaw rate < 22 deg/s) were used to exclude active turning. Third, to mitigate the effects of gravity and buoyancy, gliding events for analysis were restricted to constant depth (|depth rate| *<* 0.2 m/s and |pitch| *<* 17 deg). When the animal was moving at a constant depth, it was assumed that buoyancy and weight force were equal and opposite and did not affect the animal’s forward motion. A decision tree was employed to detect *gliding points* in the data that satisfied the specified conditions:

1. forward speed *>* 1.0 m/s
2. depth *>* 3.5 m
3. |pitch rate| *<* 20 deg/s
4. |yaw rate| *<* 22 deg/s
5. ||jerk||_1_ *<* 0.5 g/s
6. |depth rate| *<* 0.2 m/s
7. |pitch angle| *<* 17 deg

Temporally adjacent *gliding points* were grouped to form a *gliding event*. Speed measurements from a gliding event were then used with Eq. (6) to estimate the gliding drag coefficient 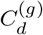, using a dual parameter sweep fitting approach presented by Zhang et al. (2020).

To characterize fluke-and-glide (FG) gait during a range of behaviors, we relaxed the restrictions on depth, pitch, and yaw rate to include more FG bouts under different conditions. Manually selected data segments composed of accelerating fluking followed by a period of gliding were included for the analysis. The start of an event was identified using the fluking signature in the data, and the endpoint of the glide was selected to equal the segment’s starting speed (with difference *<* 0.1 m/s). Fluking duration *T*_*f*_, gliding duration *T*_*g*_, duty factor *D*_*f*_ = *T*_*f*_ */*(*T*_*f*_ + *T*_*g*_), reference speed *v*_*ref*_, as well as the fluking frequency and amplitude during fluking, were subsequently parameterized from the identified FG bouts. Fluking amplitude for a stroke cycle was defined as the difference between the maximum and minimum tag measured animal pitch (i.e., peak-to-peak relative pitch difference), and the average fluking amplitude for a bout of stroke cycles was reported. The software tools used for the gait analysis were described by Zhang et al. (2021).

### Speed distribution and normalization

To investigate the speed distribution during different locomotion, we first normalize the measured speed (in m/s) by the body length of the animal to obtain a normalized speed (in l/s, i.e. body length per second). Now, let *P*_*all*_(*v*) be the probability that the animal’s speed is within the left-closed interval [*v* - 0.05, *v* + 0.05) l/s during all locomotion, then:

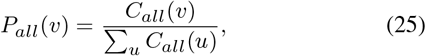

where *C*_*all*_(*v*) is the counted number of occurrences for speed that falls within the interval associated with *v* during all locomotion. Similarly, we define the probability for an animal’s reference speed (*v*_*ref*_, Eq. (1)) during fluke-and-glide (FG) gait:

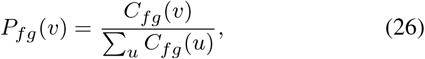

where *C*_*fg*_ (*v*) is the counted number of occurrences for speed to fall within the interval of *v* during all FG bouts. The normalized count for an FG gait bout is then defined as:

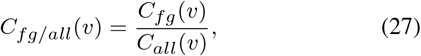

which represents the animal’s relative preference for selecting a FG gait when at speed *v*. And the associated probability for the normalized FG speed is:

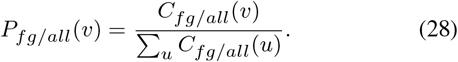

*P*_*fg/all*_(*v*) is a normalized ratio between speed distributions of FG bouts and all locomotion, making it an indicator of the animal’s tendency to select an FG gait at speed *v*.

### Statistical methods

A Linear Mixed-Effects Model (LMEM, Robinson (1991); Mclean et al. (1991); C. Pinheiro and M. Bates (1996), `fitlme` in Matlab) was employed to test the relationship between drag coefficient *C*_*d*_ and Reynolds number *Re*, under log scale, for the three participating animals. Specifically, *C*_*d*_ was the response variable, *Re* was the predictor variable, and animal identity was the grouping variable. The fixed-effects portion of the model corresponded to the *slope* and *intercept* of the global linear relationship between *C*_*d*_ and *Re* under log scale for all animals. Random-effects were associated with the (potentially correlated) *slope* and *intercept* of each animal in addition to the global trend. In total, there were two parameters for the fixed-effects and six parameters for the randomeffects assessed in the model. The *t*-statistic and *p*-value of these parameters were evaluated against a hypothesized value of 0.

The Pearson correlation coefficients (*r*-value, Fisher (1958); Kendall (1979), `corrcoef` in Matlab) together with associated *p*-values were computed to test for correlations between model estimated thrust and tag measured fluking frequency and amplitude. A plane was fitted to these three variables using robust linear regression to map from measured fluking frequency and amplitude to the estimated thrust (Fisher (1958); Huber (1981), `robustfit` in Matlab). The *p*-values and standard errors of the plane’s coefficients were assessed simultaneously. The correlation coefficient between plane predicted thrust and thrust from data points was evaluated to characterize the plane’s effectiveness in the mapping from fluking frequency and amplitude to the thrust. Robust linear regression was also used to obtain a pair-wise relationship among thrust, fluking frequency, and fluking amplitude.

## RESULTS

### Gliding drag coefficient

Data from a representative dive, along with the identified gliding events, are shown in Fig. 2 (a-d). Equation (6) was used with the gliding speed profiles (Fig. 2-e) to calculate the gliding drag coefficient 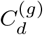. A total of 532 gliding segments with an average duration of 1.97 ± 1.09 sec (mean ± standard deviation) were identified from the animals for the analysis. The 532 pairs 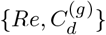 are shown in Fig. 3. A Linear Mixed-Effects Model (LMEM) was fitted to the 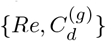 values from all animals to identify the global trend as well as assess potential individual differences. The slope and intercept of the fixed-effects (the global trend) were obtained as -0.6621 (*t*-Stat: *−* 8.0, *p*-value < 0.001) and 2.2106 (*t*-stat: 4.0, *p*-value < 0.001), yielding the following relationship between *Re* and 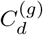:

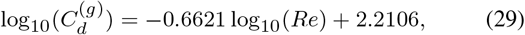

which can be rearranged to give:

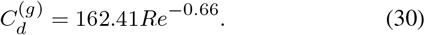

**Fig. 2.**
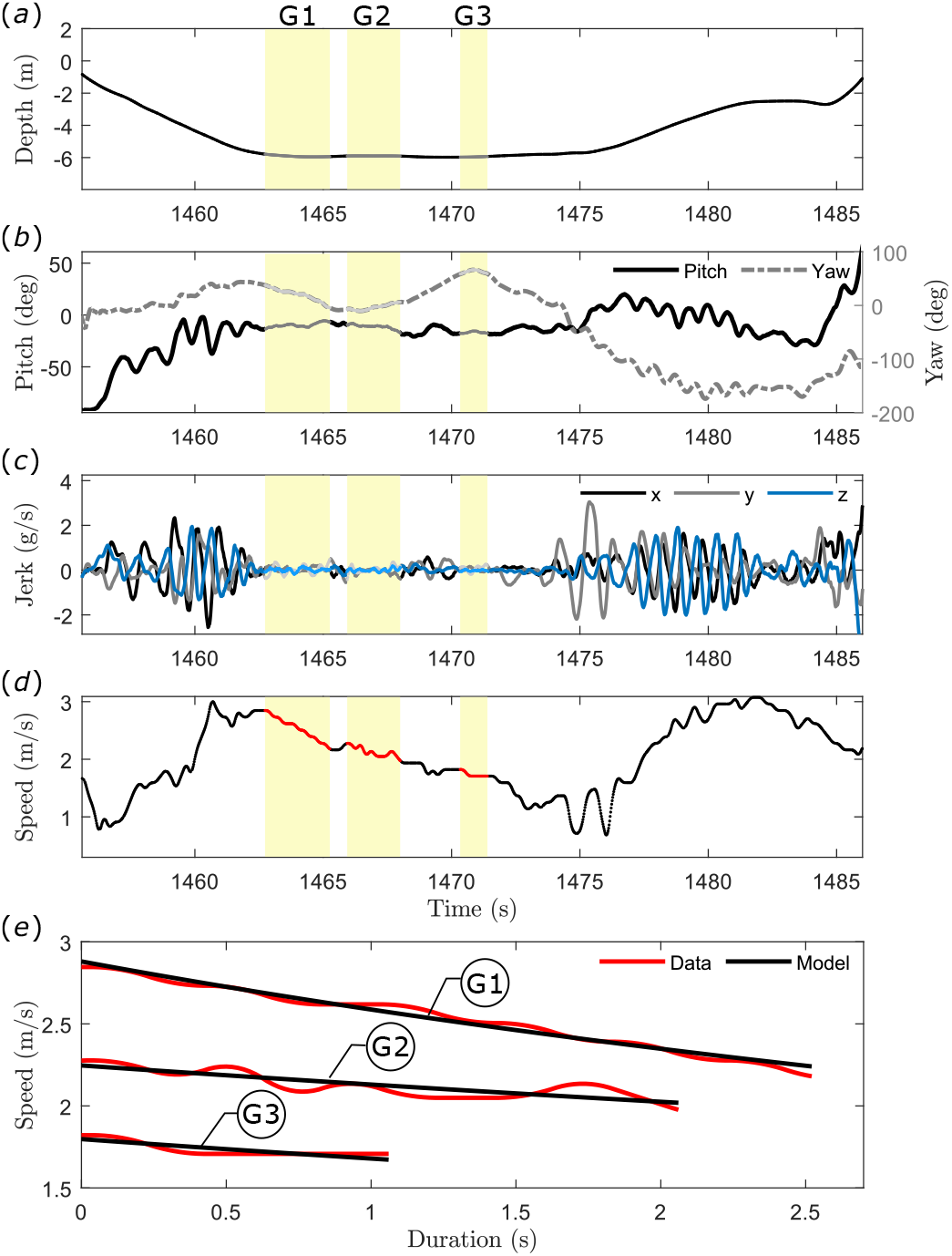
Signal features of an example section of data (a-d) and three extracted gliding events’ speed profiles (G1, G2, G3) with fitted dynamic models (e) are shown in the plots.

**Fig. 3.**
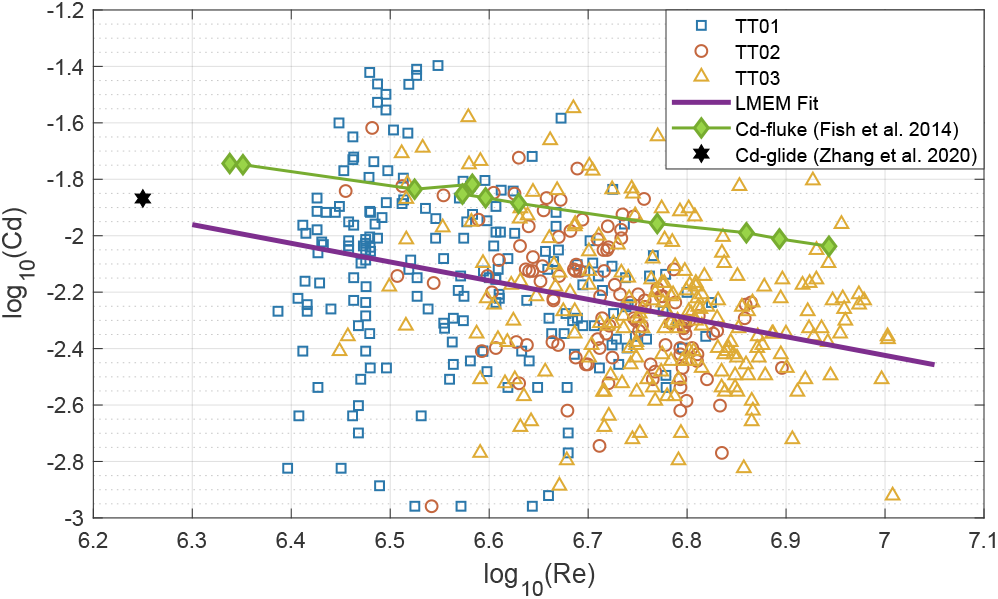
A Linear Mixed Effects Model (LMEM) fitted to the resulting 532 pairs of gliding {*Re, C*_*d*_} values from all animals (TT01, TT02, TT03) under log scale. Results from low amplitude dolphin fluking (Fish et al. (2014)) and low-speed dolphin gliding (Zhang et al. (2020)) are also shown.

This relationship was used to estimate the gliding drag coefficient 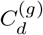 for the modeling analysis. Figure 3 presents 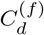 from Fish et al. (2014) 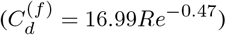, the average 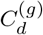 estimate from Zhang et al. (2020), and the gliding coefficients found in this work for comparison. The random-effects between *Re* and 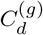 caused by individual differences were insignificant, with parameters for the random-effects that accounted for the individual differences in the LMEM all smaller than 0.001 (absolute value) and the absolute values of their 95% confidence intervals all smaller than 0.001.

### Fluke-and-glide (FG) bouts

For the modeling analysis, 783 FG bouts, with an average fluking duration *T*_*f*_ of 4.3 ± 2.1 sec and gliding duration *T*_*g*_ of 5.0 ± 2.0 sec, were identified from the data of the three animals. Duty factor, 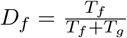, of the FG gait had a value of 0.45 ± 0.11. Figure 4 presents the relative pitch (i.e. zero centered pitch), normalized speed, estimated trust force, and estimated trust power of an example FG bout.

**Fig. 4.**
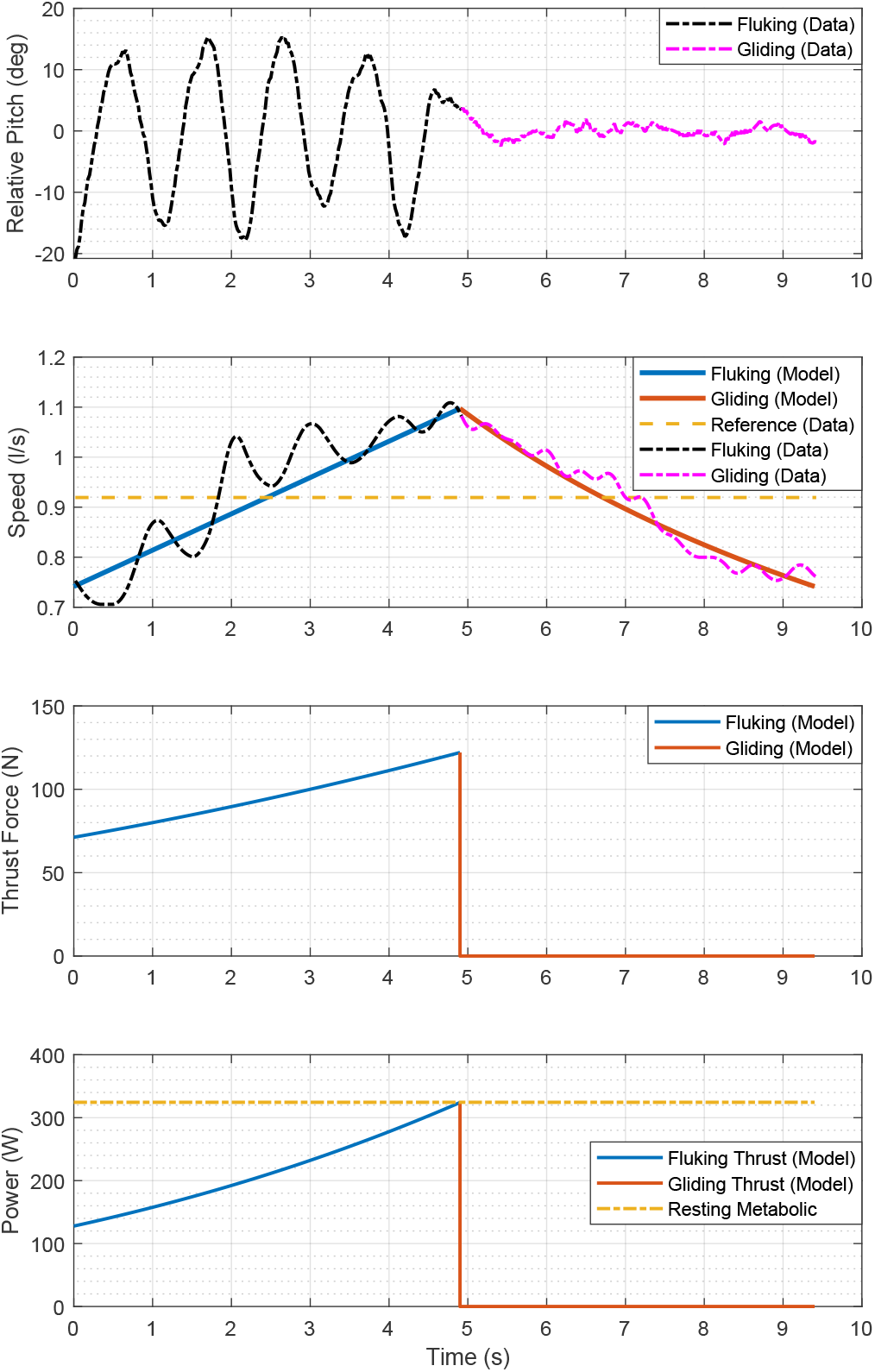
The relative pitch (i.e. zero centered pitch), speed, model estimated thrust force, and model estimated power of an example fluke-and-glide (FG) bout. The reference speed of this bout is 0.91 l/s, with fluking duration being 4.9 s, gliding duration being 4.5 s, and duty factor being 0.52. The average fluking frequency and amplitude during the fluking phase are 1.0 Hz and 27.0 degree, respectively.

The speed distributions of the animals while swimming (speed greater than 0.3 m/s) are shown in Fig. 5. Speed distributions varied between animals, with an overall swimming speed of around 0.45 l/s being most likely during all swimming bouts for all animals. FG gait was more likely to occur at slightly higher speeds (0.6 - 0.9 l/s). The normalized ratio between FG speed distribution and the general speed distribution (Eq. (28)) indicates that FG bouts were more likely to be employed by the animals at around 0.9 - 1.1 l/s (2.2 - 2.7 m/s for a body length of 2.45 m). This is a speed value near the previously reported most-efficient bottlenose dolphin swimming speed of 2.1 m/s (Williams et al. (1992)).

**Fig. 5.**
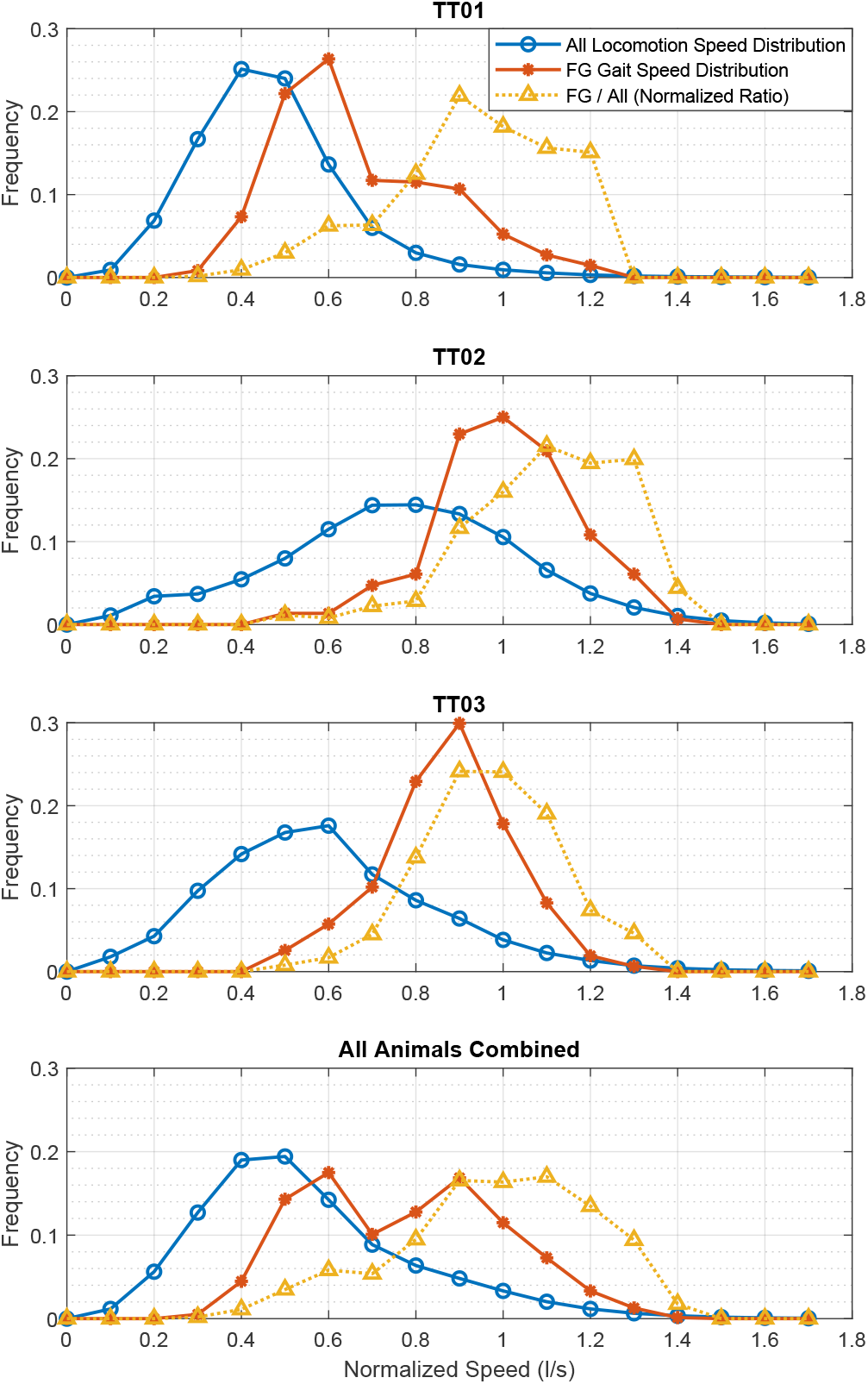
The normalized speed (unit in body length per second) distribution of all locomotion, sampled FG bouts, and the normalized ratio between FG bouts and all locomotion (Eq. (28)) from three subjects.

### Cost of transport

The COT and MECOT values associated with the 783 FG bouts were estimated and plotted against the corresponding reference speed in Fig. 6 and color-coded by duty factor. The estimated COT and MECOT of constant speed continuous fluking gait are also shown in the figures. The COT *vs*. speed relationship demonstrated a decreasing-then-increasing trend, while the MECOT *vs*. speed relationship presented a monotonically increasing trend. For a fixed speed, both COT and MECOT predicted a higher cost with increasing duty factor, where constant speed continuous fluking has the highest cost. The cost difference between continuous fluking and FG gait increases with speed. At a set of representative parameters, {*v*_*ref*_ = 1 l/s, *T*_*g*_ = 5 sec, *D*_*f*_ = 0.45}, the COT and MECOT of a FG bout were 2.53 and 0.35 J · m^*−*1^ · kg^*−*1^ respectively, corresponding to 18.9% and 27.1% reductions compared to the continuous fluking gait at the same reference speed.

**Fig. 6.**
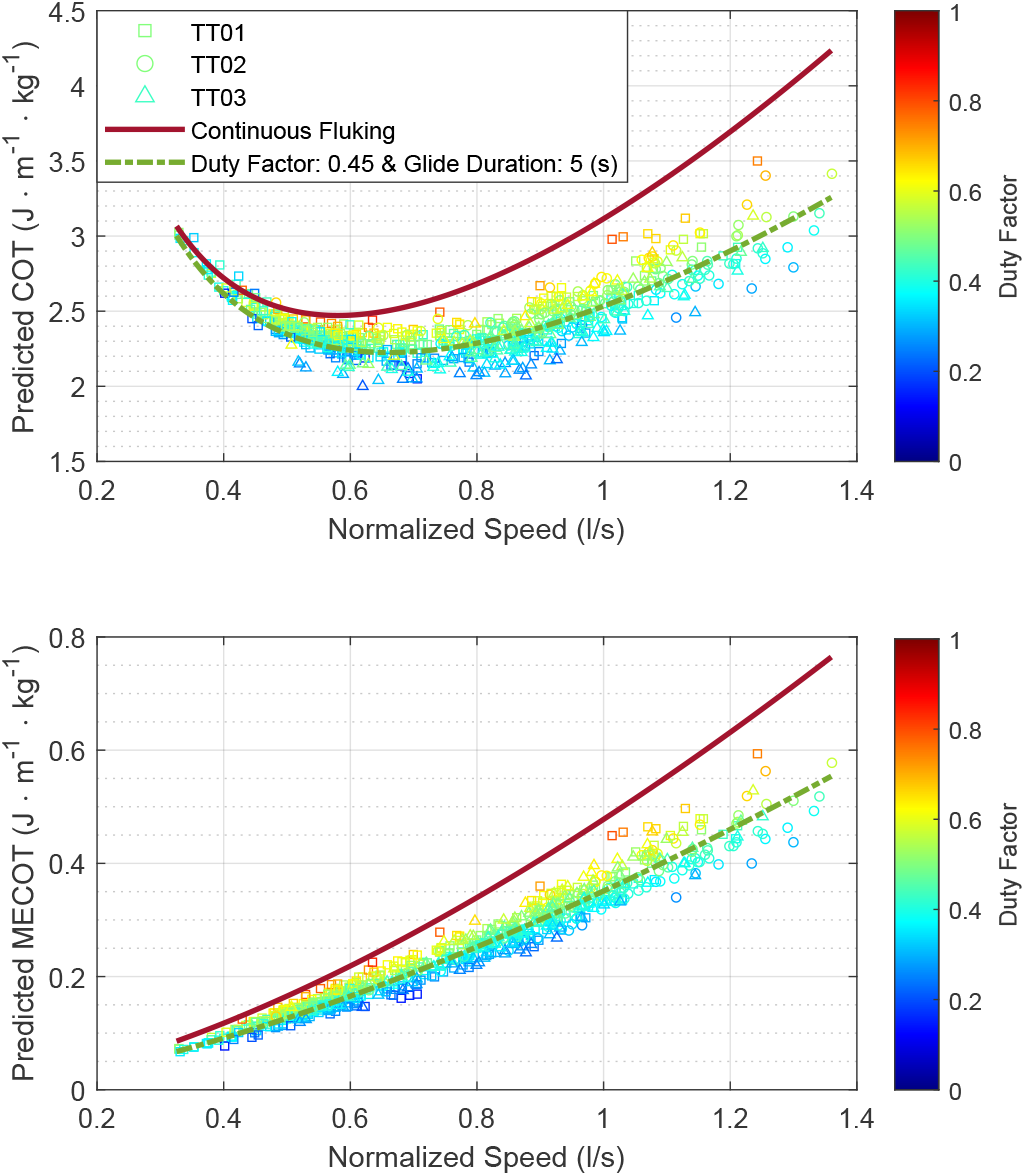
The predicted metabolic energy cost of transport (COT, top) and mechanical energy cost of transport (MECOT, bottom) of the 783 identified fluke-and-glide (FG) bouts from 3 animals plotted over the associated speeds with color codes for their duty factors. The COT and MECOT of constant speed continuous fluking gait, with an effective duty factor of 1, were compared against the FG bouts. The curves of a FG gait with duty factor equals 0.45 and gliding duration equals 5 seconds were also shown.

### Fluking dynamics

Average model estimated thrust forces (*F*_*thrust*_, Eq. (2)) during the fluking phase were normalized against the animal’s body weight (i.e. *F*_*thrust*_*/*(*mg*), unit-less, where *g* = 9.8 m/s^2^), colorcoded by duty factor, and plotted against the reference speeds in Fig. 7. In the displayed figure, the normalized thrust is positively correlated with speed (*r*-value: 0.895, *p*-value < 0.001). A gait with a higher duty factor resulted in a lower average thrust during the fluking phase at a given reference speed. The normalized thrust ranged from 0.01 to 0.15 in the identified FG bouts for these animals. Additionally, the relationships between normalized thrust and the measured fluking frequency (*Freq*, in Hertz) and amplitude (*Amp*, in degree) are presented in Fig. 8. Significant positive correlations existed between thrust and fluking frequency (*r*-value: 0.533, *p*-value < 0.001):

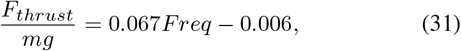

as well as between thrust and fluking amplitude (*r*-value: 0.445, *p*-value < 0.001):

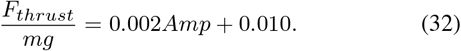

**Fig. 7.**
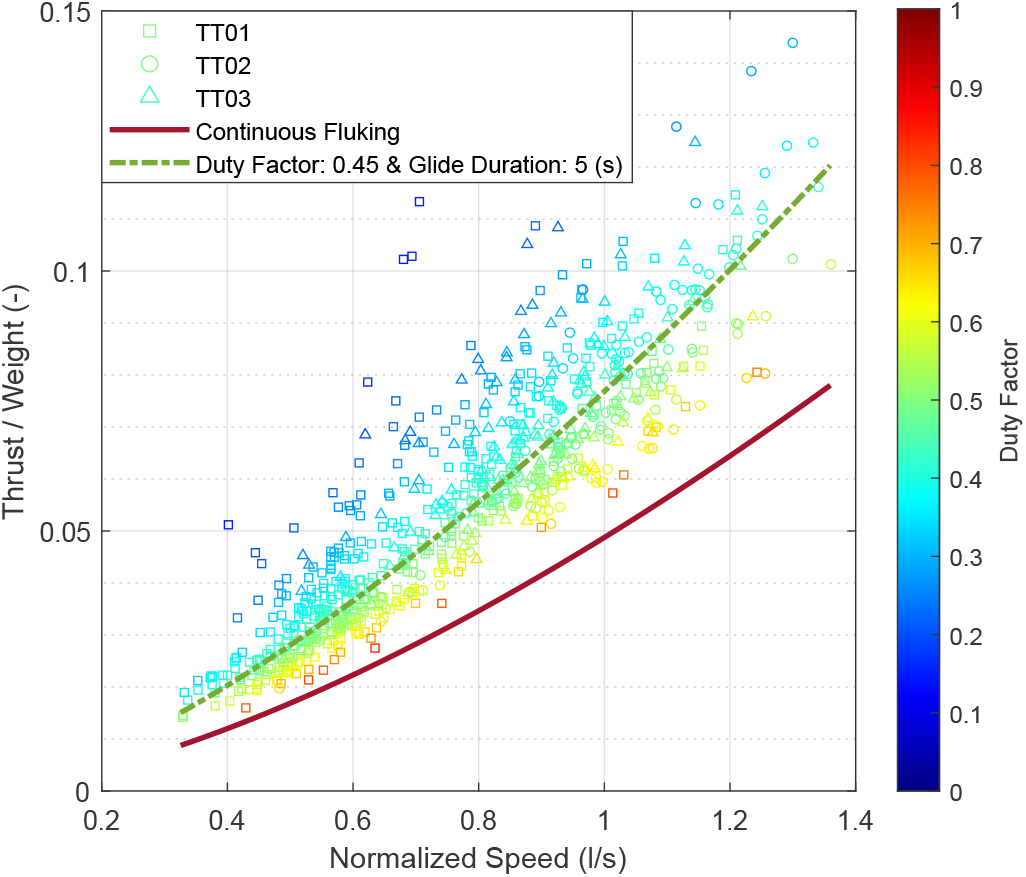
The predicted thrust (normalized over body weight) during fluking for the FG bouts and constant speed continuous fluking gait plotted over the associated speeds with color codes for their duty factors. The curve of a FG gait with duty factor equals 0.45 and gliding duration equals 5 seconds was also shown.

**Fig. 8.**
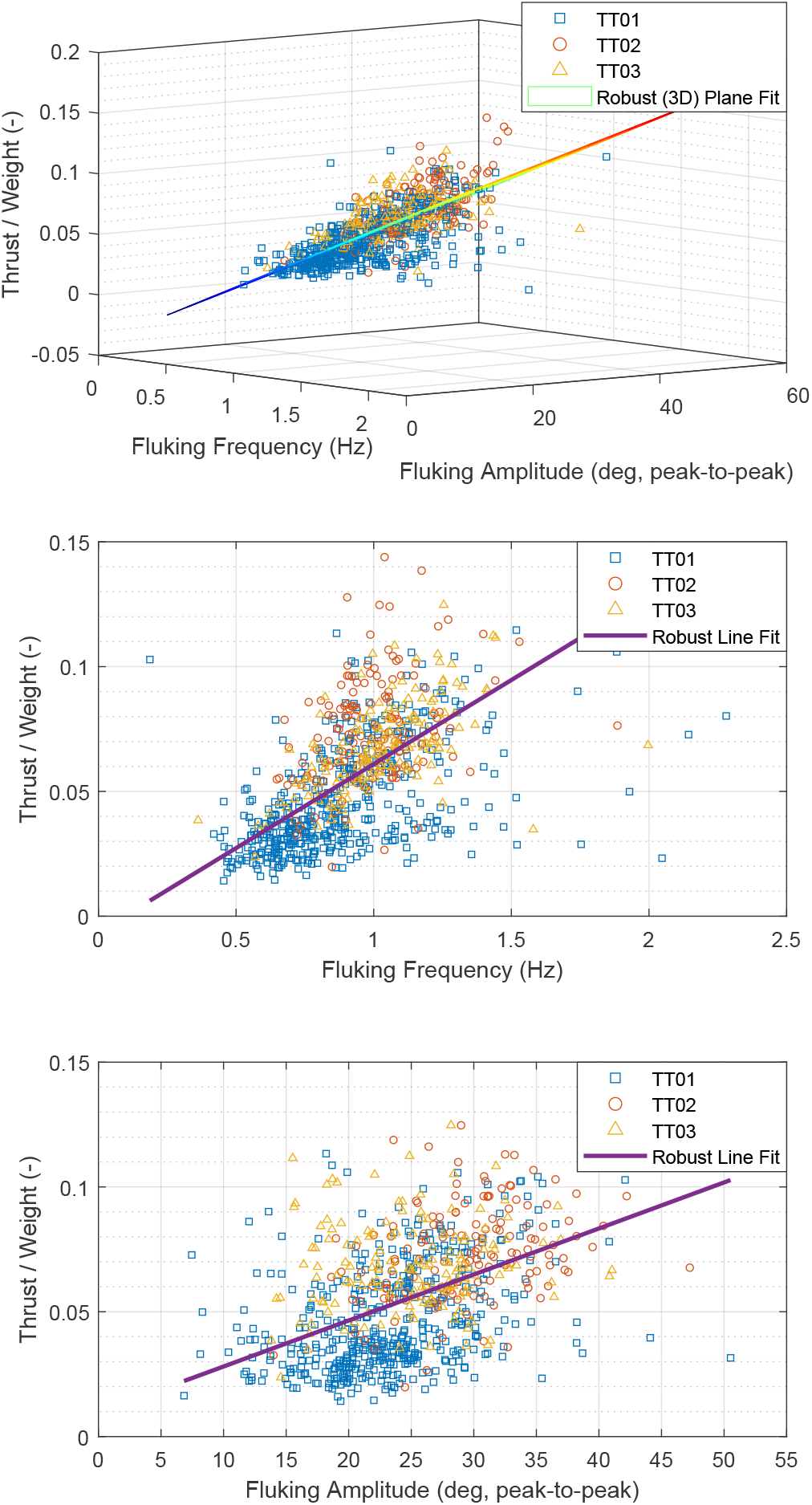
The predicted thrust (normalized over body weight) during fluking for the FG bouts was plotted against the measured fluking frequency and amplitude (peak-to-peak). A plane was robustly fitted to the data points from all three animals in 3D (top). The 2D view of the data points was also shown: thrust *vs*. fluking frequency (middle) and thrust *vs*. fluking amplitude (bottom).

The robust linear regression fit between thrust, fluking frequency, and fluking amplitude was determined to be:

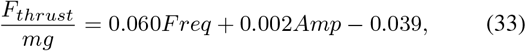

with *p*-values for the coefficients all smaller than 0.001 and standard errors being 0.002, 0.0001, and 0.003, respectively for the three coefficients. The root mean square error (RMSE) of the fit from the robust linear regression was 0.016 (unit-less). A comparison between the estimated thrust and the thrust predicted by the regression plane in a Pearson correlation test gave an *r*-value of 0.682 with a *p*-value < 0.001, which was a tighter correlation than either the thrust-frequency relationship (*r*-value: 0.533, *p*-value < 0.001) or the thrust-amplitude relationship (*r*-value: 0.445, *p*-value < 0.001). Indicating that the plane fit was better than using either frequency or amplitude alone.

For the relationship between fluking frequency and amplitude during the fluking phase of the 783 FG bouts (Fig. 9-top), no significant correlation was detected between the two parameters over the entire set of data points (*r*-value: 0.026, *p*-value: 0.460). However, correlations emerged when a narrower set of points were analyzed. The complete set of points was gathered into six equal-sized bins (131 points each) based on their normalized thrust so that points in the same bin share similar thrust magnitude. Statistical results showed that five out of the six bins presented a significant negative correlation between fluking frequency and amplitude (Fig. 9-bottom six). The negative slop of the linear relationship became steeper as the trust increased. And the strongest correlation occurred in the bin associated with the highest thrust range.

**Fig. 9.**
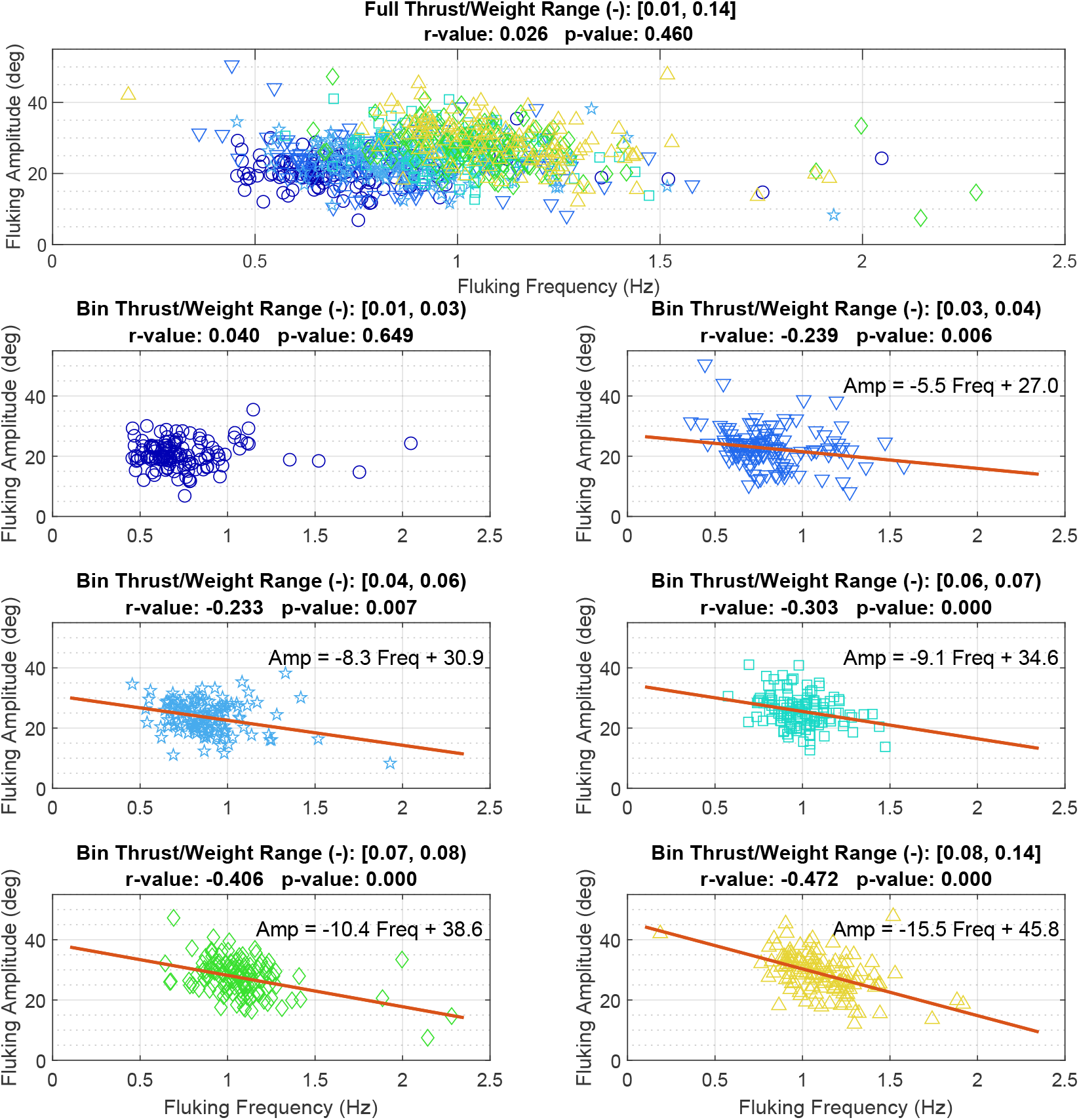
During the fluking phase of the 783 FG bouts from the three animals (top subplot, color-coded in accordance with the bottom six subplots), the fluking frequency and amplitude were gathered into six equal-sized bins (131 points each, bottom six subplots) based on their estimated thrust (normalized over body weight). The thrust range, together with the *r*-value and *p*-value for a Pearson correlation test, were indicated above each subplot. Five out of the six bins presented a significant negative correlation (*α* = 0.01) between fluking frequency and amplitude. A robust linear regression line was fitted to the data from the bin that presents significant correlation. The equation associated with the regression line was explicitly shown in the subplot.

## DISCUSSION

This work provides insight into the dynamics and energetics of a fluke-and-glide (FG) gait using physics-based models and new estimates of gliding drag coefficients of bottlenose dolphins over a range of swimming speeds. As expected, drag during gliding (Fig. 3) is lower than during low amplitude fluking (Fish et al. (2014)). The dynamic dorsal-ventral bending of the body used to generate propulsive force for locomotion results in body configurations that create more drag than the static pose used during a glide. The Bone-Lighthill boundary layer thinning hypothesis also suggests that undulating motion increases skin friction drag (Lighthill (1971); Akoz and Moored (2018)). The reduction in drag is significant across the range of speeds. For example, animal TT01 experiences 104% more drag during low amplitude fluking than during a glide at 2.25 m/s. This significant reduction contributes to the energetic benefits of FG gait observed in the swimming patterns of whales and dolphins (Shorter et al. (2017)).

Energetic benefits of FG gait were then further investigated using the proposed model and parameters identified from observed bouts of FG gait. The parabolic trend observed in the metabolic energy cost of transport (COT, Fig. 6-top) was the result of two factors. First, at lower speeds (< 0.5 l/s) the animal’s resting/baseline metabolic resulted in higher COT, since the animal still consumes metabolic energy (the numerator *E* in Eq. (18)) during small displacements (the denominator *S* in Eq. (18)). Second, the quadratic relationship between drag force and speed (Eq. (3)) creates significantly larger COT at higher speed as the animal has to increase thrust production to overcome the drag. While COT is an estimate of the overall metabolic energy cost of the animal during locomotion, the mechanical energy cost of transport (MECOT) describes only the amount of mechanical energy required for locomotion. The monotonic trend in the MECOT *vs*. speed relationship (Fig. 6-bottom) is mainly a result of the quadratic relationship between drag force and speed. MECOT better captures the interaction between the animal, as one dynamical system, and the environment.

Despite the differences between COT and MECOT, both metrics indicate that an FG gait is more efficient than a corresponding constant speed continuous fluking gait. Results from the model at a 1.0 l/s swimming speed with a representative set of gait parameters show a 18.9% reduction in the COT and a 27.1% reduction in the MECOT compared to constant speed fluking. A 18.9% savings for bottlenose dolphins are smaller than the over 50% estimated savings for a fish (Weihs (1974)). Still, for dolphins that may travel more than 50 km per day (Mate et al. (1995)), a 18.9% cost savings would result in a significant reduction in the foraging effort required to maintain the animal’s energy budget. Further, the energetic benefits of FG gait grow as swimming speed increases (Fig. 6), supporting the observation that the animals may preferentially select an FG gait at higher speeds (Fig. 5).

Additional results in Fig. 6 demonstrate that COT and MECOT of FG gait are lower when the duty factor is lower. That is, swimming efficiency improves when the animal spends more time gliding and less time fluking. While a small duty factor with as much gliding as possible leads to improved COT and MECOT, a low duty factor also requires higher acceleration, thus higher thrust force (Fig. 7), during the limited amount of fluking to build up speed for the glide. A higher thrust requirement leads to two caveats. First, the metabolic cost to support the burst could be higher than estimated. And second, the animal may have to increase its fluking amplitude to satisfy the dynamic demand of higher thrust.

To further investigate the dynamical properties during the fluking phase of the FG bouts, the relationships between average thrust force, fluking frequency, and fluking amplitude were investigated (Fig. 8, Eq. (33)). It was observed that thrust is positively correlated with fluking frequency and amplitude. Previous studies (Fish (1998); Rohr and Fish (2004)) have investigated the relationship between the steady state swimming speed of cetaceans and their fluking frequency and amplitude, and found a positive correlation between swimming speed and fluking frequency, yet an insignificant correlation between speed and amplitude. The data presented here builds on these previous observations of gait kinematics during steady-state swimming (i.e., near-constant speed) by including transient state dynamics as the animals accelerate before a glide. A negative correlation was found between fluking frequency and amplitude within a given range of thrust force (Fig. 9), which expands the observation that frequency and amplitude were not correlated (Rohr and Fish (2004)) when all data were shown together. These results indicate that the animals may modulate both frequency and amplitude to generate a given thrust force and higher average thrust force demands increased fluking frequency and amplitude. Such an increased rate of force development would likely lead to extra metabolic cost in dolphins, as has been described in the human biomechanics literature (van der Zee and Kuo (2021)), due to the additional active calcium transport in muscle tissues. Further, it has been shown that maintaining a higher force (without doing any mechanical work) also costs more metabolic energy in dolphins (Williams et al. (1993)), which suggests that employing a higher muscle force induces additional metabolic cost. As such, the energy conversion ratios, *η*_*mp*_*η*_*mm*_ in Eq. 19, should be investigated in future work to improve the physics-based metabolic energy estimates.

Another consideration regarding the model is that the drag coefficient for fluking 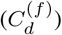 in this work comes from estimates made by Fish et al. (2014) during low amplitude dolphin fluking. We speculate that the drag coefficient associated with larger amplitude fluking would be higher. Thus making the actual mechanical work higher than estimated in this work when large amplitude motion was used to generate thrust. Further investigation could be conducted into this matter as well.

The benefits of the physics-based data-driven approach in this work were threefold: first, there was no need for dedicated experiments that involved divers or underwater cameras for drag estimation. Second, data were collected during the animal’s daily routine, linking the theoretical model with the animals’ self-selected behavior. Third, the many hours of tag data resulted in a large number of events for the analysis, helping to reduce the noise and uncertainty in the estimated parameters. While expert knowledge was used to select features from the sensor data to create a decision tree classifier, and to identify the FG bouts from data, other automated machine learning algorithms could be used to identify events of interest in the data (Kabra et al. (2013); Zhang et al. (2018); Sibal et al. (2021); Zhang (2021)). These automated approaches could further reduce the expert knowledge required for signal feature selection and event detection for potentially more complicated movement patterns. However, they would still need a labeled data set to train the algorithms. As an extension of this work, the on-body tag measurements could be combined with a model of the animal’s full body kinematics to provide thrust and drag estimates under different scenarios, providing more detailed insights about the locomotion of these animals. Future work should also use tag data from dolphins in the wild to investigate the cost of transport associated with swimming gaits used in a different environmental context.

Overall, this work presented a data-driven physics-based approach to estimate gliding drag coefficients, energetic efficiency, and gait dynamics of bottlenose dolphins during horizontal intermittent locomotion. The COT and MECOT were investigated together with gait kinematics during the bursts of accelerating speed generated for the FG gait. The results indicate that horizontal fluke-and-glide gait, in addition to porpoising/leaping (Au and Weihs (1980); Hui (1989); Fish and Hui (1991)), wave riding (Williams et al. (1992)), drafting (Weihs (2004)), and gliding during diving (Williams (2001); Williams et al. (2000); Skrovan et al. (1999)), is a behavioral mechanism that can enhance the locomotion efficiency of swimming dolphins.

## Acknowledgements

We would like to thank the Seven Seas animal care specialists at the Brookfield Zoo who facilitated this study and the animals in their care. We would also like to thank Sarah Breen-Bartecki and Bill Zeigler for their continued support. The study protocol was approved by the Chicago Zoological Society IACUC and the University of Michigan Animal Welfare Committee (IACUC, #PRO00008825).

## Data availability

Data and code are available upon request and approval.

## REFERENCES

Akoz, E. and Moored, K. W. (2018). Unsteady propulsion by an intermittent swimming gait. Journal of Fluid Mechanics 834, 149–172.

Ashraf, I., van Wassenbergh, S. and Verma, S. (2020). Burst-and-coast swimming is not always energetically beneficial in fish (Hemigrammus bleheri). Bioinspiration and Biomimetics 16, 016002.

Au, D. and Weihs, D. (1980). At high speeds dolphins save energy by leaping. Nature 284, 548–550.

Bilo, D. and Nachtigall, W. (1980). A Simple Method to Determine Drag Coefficients in Aquatic Animals. Journal of Experimental Biology 87, 357–359.

C. Pinheiro J. and M. Bates D. (1996). Unconstrained parametrizations for variance-covariance matrices. Statistics and Computing 6, 289–296.

Conti, W. (2021). Edge Innovations: https://www.edgefx.com/.

Faraji, S., Wu, A. R. and Ijspeert, A. J. (2018). A simple model of mechanical effects to estimate metabolic cost of human walking. Scientific Reports 8, 1–12.

Feldkamp, S. D. (1987). Swimming in the California sea lion: morphometrics, drag and energetics. The Journal of Experimental Biology 131, 117–135.

Fish, F. E. (1993). Power Output and Propulsive Efficiency of Swimming Bottlenose Dolphins (Tursiops Truncatus). Journal of Experimental Biology 185, 179–193.

Fish, F. E. (1998). Comparative kinematics and hydrodynamics of odontocete cetaceans: morphological and ecological correlates with swimming performance. Journal of Experimental Biology 201, 2867–2877.

Fish, F. E., Fegely, J. F. and Xanthopoulos, C. J. (1991). Burst-and-coast swimming in schooling fish (Notemigonus crysoleucas) with implications for energy economy. Comparative Biochemistry and Physiology – Part A: Physiology 100, 633–637.

Fish, F. E. and Hui, C. A. (1991). Dolphin swimming-a review. Mammal Review 21, 181–195.

Fish, F. E., Legac, P., Williams, T. M. and Wei, T. (2014). Measurement of hydrodynamic force generation by swimming dolphins using bubble DPIV. Journal of Experimental Biology 217, 252–260.

Fisher, R. (1958). Statistical Methods for Research Workers. Hafne, 13 edition.

Gabaldon, J., Anderson, E. J., Barton, K., Shorter, K. A., Johnson-Roberson, M., Turner, E. L. and Johnson, M. (2019). Integration, calibration, and experimental verification of a speed sensor for swimming animals. IEEE Sensors Journal PP, 1–1.

Gabaldon, J., Zhang, D., Rocho-Levine, J., Moore, M., van der Hoop, J., Barton, K. and Shorter, K. A. (2022). Tag-based estimates of bottlenose dolphin swimming biomechanics and energetics. Submitted for Publication.

Gleiss, A. C., Jorgensen, S. J., Liebsch, N., Sala, J. E., Norman, B., Hays, G. C., Quintana, F., Grundy, E., Campagna, C., Trites, A. W. et al. (2011). Convergent evolution in locomotory patterns of flying and swimming animals. Nature Communications 2.

Hertel, H. (1966). Structure, form, movement. New York: Reinhold Publishing Co.

Huber, P. J. (1981). Robust Statistics. Wiley.

Hui, C. A. (1989). Surfacing Behavior and Ventilation in Free-Ranging Dolphins. Journal of Mammalogy 70, 833–835.

Kabra, M., Robie, A. A., Rivera-Alba, M., Branson, S. and Branson, K. (2013). JAABA: Interactive machine learning for automatic annotation of animal behavior. Nature Methods 10, 64–67.

Kendall, M. (1979). The Advanced Theory of Statistics. Macmillan.

Kramer, D. L. and McLaughlin, R. L. (2001). The behavioral ecology of intermittent locomotion. American Zoologist 41, 137–153.

Lang, T. G. (1975). Speed, Power, and Drag Measurements of Dolphins and Porpoises. In Swimming and Flying in Nature, pp. 553–572. New York: Springer.

Lighthill, M. J. (1971). Large-amplitude elongated-body theory of fish locomotion. Proceedings of the Royal Society of London. Series B. Biological Sciences 179, 125–138.

Madgwick, S. O., Harrison, A. J. and Vaidyanathan, R. (2011). Estimation of IMU and MARG orientation using a gradient descent algorithm. In IEEE International Conference on Rehabilitation Robotics, pp. 1–7. IEEE.

Mate, B. R., Kelly A. Rossbach, Sharon L. Nieukirk, Wells, R. S., Irvine, A. B., Scott, M. D. and Read, A. J. (1995). Satellite-monitored movements and dive behavior of a bottlenose dolphin (Tursiops truncatus) in Tampa Bay, Florida. Marine Mammal Science 11, 452–463.

McHenry, M. J. and Lauder, G. V. (2005). The mechanical scaling of coasting in zebrafish (Danio rerio). Journal of Experimental Biology 208, 2289–2301.

Mclean, R. A., Sanders, W. L. and Stroup, W. W. (1991). A Unified Approach to Mixed Linear Models. The American Statistician. American Statistical Association 45, 54–64.

Miller, P. J., Johnson, M. P., Tyack, P. L. and Terray, E. A. (2004). Swimming gaits, passive drag and buoyancy of diving sperm whales Physeter macrocephalus. Journal of Experimental Biology 207, 1953–1967.

Müller, U. K., Stamhuis, E. J. and Videler, J. J. (2000). Hydrodynamics of unsteady fish swimming and the effects of body size: Comparing the flow fields of fish larvae and adults. Journal of Experimental Biology 203, 193–206.

Noren, S. R., Redfern, J. V. and Edwards, E. F. (2011). Pregnancy is a drag: Hydrodynamics, kinematics and performance in pre-and postparturition bottlenose dolphins (Tursiops truncatus). Journal of Experimental Biology 214, 4151–4159.

Purves, P. E., Dudok van Heel, W. H. and Jonk, A. (1975). Locomotion in dolphins Part I: Hydrodynamic experiments on a model of the bottle-nosed dolphin, Tursiops truncatus, (Mont.).

Rayner, J. M., Viscardi, P. W., Ward, S. and Speakman, J. R. (2001). Aerodynamics and energetics of intermittent flight in birds. American Zoologist 41, 188–204.

Robinson, G. K. (1991). That BLUP Is a Good Thing: The Estimation of Random Effects. Statistical Science 6, 15–32.

Rohr, J. J. and Fish, F. E. (2004). Strouhal numbers and optimization of swimming by odontocete cetaceans. Journal of Experimental Biology 207, 1633–1642.

Sachs, G. (2015). New model of flap-gliding flight. Journal of Theoretical Biology 377, 110–116.

Schmidt-Nielsen, K. (1972). Locomotion: Energy cost of swimming, flying, and running. Science 177, 222–228.

Shorter, K. A., Shao, Y., Ojeda, L., Barton, K., Rocho-Levine, J., van der Hoop, J. and Moore, M. (2017). A day in the life of a dolphin: Using biologging tags for improved animal health and well-being. Marine Mammal Science 33, 785–802.

Sibal, R., Zhang, D., Rocho-Levine, J., Shorter, K. A. and Barton, K. (2021). Bidirectional LSTM Recurrent Neural Network Plus Hidden Markov Model for Wearable Sensor-Based Dynamic State Estimation. ASME Letters in Dynamic Systems and Control 1, (021002)1–5.

Skrovan, R., Williams, T. M., Berry, P. S., Moore, P. W. and Davis, R. W. (1999). The Diving Physiology of Bottlenose Dolphins. Journal of Experimental Biology 202, 2749–2761.

Stelle, L. L., Blake, R. W. and Trites, A. W. (2000). Hydrodynamic drag in Steller sea lions (Eumetopias jubatus). Journal of Experimental Biology 203, 1915–1923.

Tobalske, B. W. (2001). Morphology, velocity, and intermittent flight in birds. American Zoologist 41, 177–187.

van der Hoop, J. M., Fahlman, A., Hurst, T., Rocho-Levine, J., Shorter, K. A., Petrov, V. and Moore, M. J. (2014). Bottlenose dolphins modify behavior to reduce metabolic effect of tag attachment. Journal of Experimental Biology 217, 4229–4236.

van der Hoop, J. M., Fahlman, A., Shorter, K. A., Gabaldon, J., Rocho-Levine, J., Petrov, V. and Moore, M. J. (2018). Swimming energy economy in bottlenose dolphins under variable drag loading. Frontiers in Marine Science 5, 465.

van der Zee, T. J. and Kuo, A. D. (2021). The high energetic cost of rapid force development in muscle. The Journal of experimental biology 224.

Videler, J. J. and Weihs, D. (1982). Energetic advantages of burst-and-coast swimming of fish at high speeds. Journal of Experimental Biology 97, 169–178.

Vogel, S. (1996). Life in Moving Fluids: The Physical Biology of Flow. Princeton University Press, 2nd revise edition.

Watanuki, Y., Takahashi, A., Daunt, F., Wanless, S., Harris, M., Sato, K. and Naito, Y. (2005). Regulation of stroke and glide in a foot-propelled avian diver. Journal of Experimental Biology 208, 2207–2216.

Weihs, D. (1973). Mechanically efficient swimming techniques for fish with negative buoyancy. Journal of Marine Research 31, 194–209.

Weihs, D. (1974). Energetic advantages of burst swimming of fish. Journal of Theoretical Biology 48, 215–229.

Weihs, D. (2004). The hydrodynamics of dolphin drafting. Journal of Biology 3, 1–16.

Weimerskirch, H., Martin, J., Clerquin, Y., Alexandre, P. and Jiraskova, S. (2001). Energy saving in flight formation. Nature 413, 697–698.

Williams, T. M. (2001). Intermittent swimming by mammals: A strategy for increasing energetic efficiency during diving. American Zoologist 41, 166–176.

Williams, T. M., Davis, R. W., Fuiman, L. A., Francis, J., Le Boeuf, B. J., Horning, M., Calambokidis, J. and Croll, D. A. (2000). Sink or swim: Strategies for cost-efficient diving by marine mammals. Science 288, 133–136.

Williams, T. M., Friedl, W. A., Fong, M. L., Yamada, R. M., Sedivy, P. and Haun, J. E. (1992). Travel at low energetic cost by swimming and wave-riding bottlenose dolphins. Nature 355, 821–823.

Williams, T. M., Friedl, W. A. and Haun, J. E. (1993). The physiology of bottlenose dolphins (Tursiops truncatus): heart rate, metabolic rate and plasma lactate concentration during exercise. Journal of experimental biology 179, 31–46.

Williams, T. M., Kendall, T. L., Richter, B. P., Ribeiro-French, C. R., John, J. S., Odell, K. L., Losch, B. A., Feuerbach, D. A. and Stamper, M. A. (2017). Swimming and diving energetics in dolphins: A stroke-by-stroke analysis for predicting the cost of flight responses in wild odontocetes. Journal of Experimental Biology 220, 1135–1145.

Wu, Z., Yu, J., Yuan, J. and Tan, M. (2019). Towards a gliding robotic dolphin: Design, modeling, and experiments. IEEE/ASME Transactions on Mechatronics 24, 260–270.

Yazdi, P., Kilian, A. and Culik, B. M. (1999). Energy expenditure of swimming bottlenose dolphins (Tursiops truncatus). Marine Biology 134, 601–607.

Zhang, D. (2021). From AI to IA : Towards Intelligent Analysis of Cooperative Behavior in Bottlenose Dolphins. Ph.D. thesis, University of Michigan, Ann Arbor.

Zhang, D., Goodbar, K., West, N., Lesage, V., Parks, S., Wiley, D., Barton, K. and Shorter, K. A. (2021). Pose-gait analysis for cetaceans with biologging tag. bioRxiv pp. 1–23.

Zhang, D., Shorter, K. A., Rocho-Levine, J., van der Hoop, J., Moore, M. and Barton, K. (2018). Behavior Inference From Bio-Logging Sensors: A Systematic Approach for Feature Generation, Selection and State Classification. In Proceedings of the ASME 2018 Dynamic Systems and Control Conference, pp. 1–10. September 30–October 3, 2018, Atlanta, Georgia, USA: ASME.

Zhang, D., van der Hoop, J. M., Petrov, V., Rocho-Levine, J., Moore, M. J. and Shorter, K. A. (2020). Simulated and experimental estimates of hydrodynamic drag from bio-logging tags. Marine Mammal Science 36, 136–157.

